# Lyme Disease IgG N-linked Glycans Contrast the Canonical Inflammatory Signature

**DOI:** 10.1101/2022.05.09.491121

**Authors:** Benjamin Samuel Haslund-Gourley, Stéphane Grauzam, Anand S. Mehta, Brian Wigdahl, Mary Ann Comunale

**Affiliations:** Department of Microbiology and Immunology, Drexel University College of Medicine, Philadelphia, PA, USA; Institute for Molecular Medicine and Infectious Disease, Drexel University College of Medicine, Philadelphia, PA, USA; GlycoPath, LLC Charleston, SC; Department of Cell and Molecular Pharmacology, Medical University of South Carolina (MUSC), 173 Ashley Avenue, Charleston, SC

## Abstract

Lyme disease (LD) infection is caused by *Borrelia burgdorferi* sensu *lato*. Due to the limited presence of this pathogen in the bloodstream in humans, diagnosis of LD relies on seroconversion. Immunoglobulins produced in response to infection are differentially glycosylated to promote or inhibit downstream inflammatory responses by the immune system. IgG N-glycan responses to LD have not been characterized. In this study, we analyzed IgG N-glycans from cohorts of healthy controls, acute LD patient serum, and serum collected after acute LD patients completed a 2- to 3-week course of antibiotics and convalesced for 70-90 days. Results indicate that during the acute phase of Bb infection, IgG shifts its glycosylation profile to include structures that are not associated with the classic proinflammatory IgG N-glycan signature. This unexpected result is in direct contrast to what is reported for other inflammatory diseases. Furthermore, IgG N-glycans detected during acute LD infection discriminated between control, acute, and treated cohorts with a sensitivity of 75-100% and specificity of 94.7-100%.

**Author summary:** The causative agent of Lyme disease (LD), Borrelia burgdorferi sensu lato (Bb), is transmitted from an infected Ixodes tick into the human host dermis during the tick’s blood meal. Currently, LD is the most prevalent vector-borne disease in the US, with an estimated 476,000 annual cases. LD diagnostics rely on patient seroconversion against Bb antigens, and these tests cannot distinguish between an acute patient compared to a patient previously treated for LD. With the goal of identifying novel biomarkers associated specifically with LD infections, we analyzed the glycoprotein Immunoglobulin G (IgG) N-glycan signatures from healthy control, acute LD, and a second time point composed of the same LD patients after antibiotic therapy. We found acute LD IgG N-glycan signatures were significantly different from the canonical pro-inflammatory profile associated with most inflammatory diseases. The dramatic shifts observed in the acute LD time point were further altered at the treated time point. IgG N-glycan signature data was employed to discriminate between acute LD and healthy controls. In addition, IgG N-glycan signatures distinguished patients who completed antibiotic therapy from the acute LD timepoint. Our study will contribute to the accurate and prompt treatment of LD patients and reveals a new research avenue of immune dysregulation associated with LD.

**Graphical Abstract:** 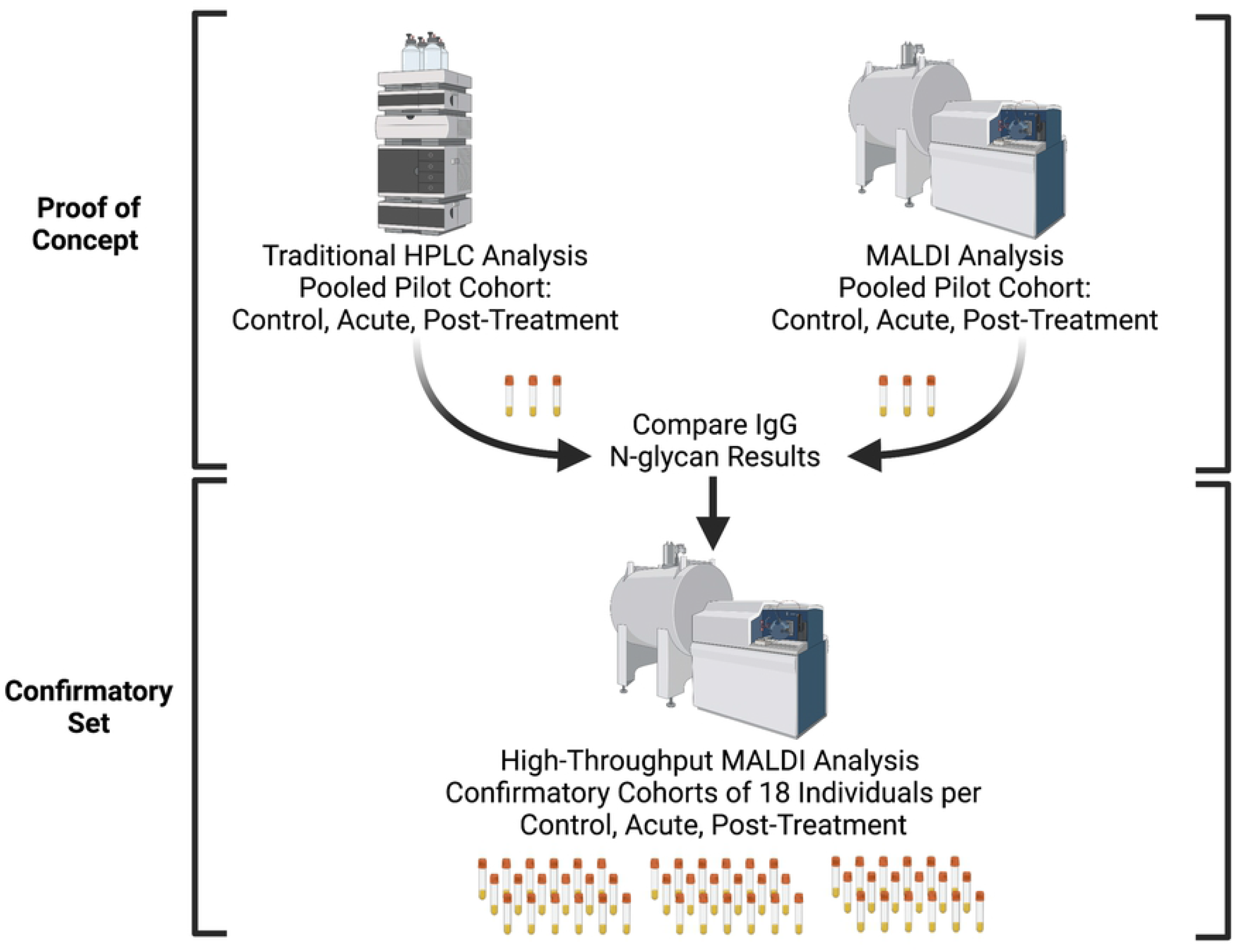

## Introduction

In 1975, a cluster of children and adults in the small community of Lyme, Connecticut experienced unusual arthritic symptoms. Today, Lyme disease (LD) is the most prevalent vector-borne disease in the US, with an estimated 476,000 annual cases [1]. The causative agent, *Borrelia burgdorferi* sensu lato (Bb), is a spirochete transmitted from the *Ixodes* tick into the human host dermis during its blood meal [2]. The spirochete leaves the blood and disseminates into multiple organ systems in as little as two weeks post-infection [3–5]. Disseminated LD is more challenging to diagnose, and delayed treatment can lead to long-term disability or death [6, 7]. The disease is endemic in the Northeastern US, and incidence rates continue to rise. The US’s annual treatment and diagnostic cost is over 4.8 billion USD [8].

Antibiotic treatment for LD in the acute phase is often curative [9–12]. However, untreated patients and a subset of treated patients progress to disseminated disease [13, 14], Disseminated disease can result in facial nerve palsy [15], Lyme Carditis [16], Lyme Arthritis [17], Lyme Neuroborreliosis [18, 19], and long-term disability [20, 21]. Persistent symptoms are reported by 10-20% of patients diagnosed and treated during the acute phase of LD. Persistent symptoms include joint pain, fatigue, and neurocognitive deficits [10, 22]. This highlights the need for an accurate early diagnosis and the ability to track disease resolution.

Current diagnosis is complicated because testing relies on indirect methods. Direct PCR and blood culture methods often fail due to the spirochete’s limited presence in the bloodstream, low bacterial counts in circulation, slow replication cycle, the requirement for complex growth media, and specialized microscopy requirements [23, 24]. Hence, indirect methods that rely on the patient’s serological response are the principal method of confirming an infection [25]. These indirect methods of acute LD diagnostics based on ELISA and western immunoblot technologies suffer from low sensitivity and a high false-positive rate. Thus, while advances in LD detection research are being made [26–30], clinicians currently lack a sensitive method to diagnose early disease. Furthermore, clinical assays are unable to determine treatment efficacy, track disease resolution [31] or diagnose subsequent infections.

Serum protein glycosylation is often altered during inflammatory and autoimmune diseases. The glycosylation profile of immunoglobulins is dynamic and offers a novel immunologic insight into the host’s response [32–35]. Glycosylation is the most abundant complex post-translational serum protein modification [36] and plays a significant role in protein structure and function *in vivo* [37, 38]. IgG has a well-characterized N-glycosylation site on the constant fragment (Fc- Asn-297) region [39]. This site contains complex biantennary glycans with varying degrees of galactose, bisecting N-acetyl-glucosamine, and sialic acid residues. Most notably, the glycans are highly core-fucosylated. IgG glycosylation is dynamic and the glycans present can affect the binding avidity to various Fcγ receptors, rate of complement activation, and release of cytokines [40–43]. Previous studies have profiled IgG N-glycans in sera obtained from patients with inflammatory disease (**Table 1)**. Reports indicate a trend for IgG N-glycan reduction in terminal galactose and sialic acid content during inflammatory diseases and this glycan signature is linked to a pro-inflammatory phenotype [44–50].

**Table 1.** Altered IgG N-glycan Profiles Detected in Inflammatory Diseases. Inflammatory diseases have serum IgG with N-glycan alterations that promote a pro-inflammatory signaling cascade due to a reduction in terminal galactose and sialic acid.

| Disease | IgG Fc N-Glycan Profile | Method of Detection | Source |
| --- | --- | --- | --- |
| Rheumatoid Arthritis (RA) and Osteoarthritis (OA) | Decreased Gal (FA2G1, FA2G2)<br>Decreased Bisects (FA2BG1, FA2BG2) | MALDI-TOF MS | [33, 34] |
| Tuberculosis | Decreased Galactose | Radiolabels and sequential exoglycosidase digestion | [32, 51] |
| Infective endocarditis | Decreased Galactose | Lectin Analysis | [52] |
| Visceral leishmaniasis | Decreased Galactose<br>Decreased Sialylation | MALDI-TOF MS | [53] |
| Hepatitis B: chronic infection | Decreased Galactose | LC-MS | [54] |
| Hepatitis C – anti-Gal IgG | Decreased Galactose<br>Increased Fucose | HPLC and Lectin | [55] |
| Liver Fibrosis | Increased Bisects<br>Decreased Galactose | MALDI Imaging | [56] |
| Inflammatory bowel disease & Crohn's disease | Decreased Galactose<br>Decreased Sialylation | UPLC | [35] |
| Endometriosis | Decreased Sialylation & Galactose<br>Increased Bisecting N-acetyl- | UPLC | [57] |
| Systemic Lupus | Decreased Galactose<br>Decreased Sialylation | UPLC | [58] |
| Moderate COVID-19 | Decreased Galactose<br>Increased Agalactose | Nano-LC-ESI-MS | [59] |

Scientists are beginning to understand how glycosylation reflects health status, and influences protein structure and function. IgG glycosylation moieties are associated with specific functions. In general, loss of galactose and sialic acid residues are reported in inflammatory states (**Table 1**), and conversely, increased galactose is associated with an anti-inflammatory state. Hence, to improve immunotherapy outcomes, pharmaceutical companies are glycoengineering monoclonal antibodies used to treat cancers and chronic diseases. For example, the effectiveness of intravenous immunoglobulin (IVIG) therapy for autoimmune conditions including Gillian-Barre, Immune Thrombocytopenic Purpura, or Kawasaki disease patients is associated with the increased sialic acid content on the Fc N-glycans of the IVIG [60–62]. In addition, therapeutic monoclonal antibodies are glycoengineered to contain specific a-fucosylated, agalactosylated N-glycans to promote superior half-life and treatment efficacy [63–66]. IgG N-glycan modulation of the immune response during disease reveals another layer of physiologic crosstalk. Evidence indicates that the repertoire of N-glycans present on IgG produced in response to vaccines is dependent on many factors, including age, inflammatory state, and the type of adjuvant [67].

It is well accepted that N-glycans modulate the function of antibodies and are altered in disease states [68]. We hypothesize that the IgG N-glycan profile of LD patients will reflect the immunological response to the acute LD infection. Thus, this first report of total IgG N-glycans associated with LD will provide insight into the inability of the host immune system to resolve Bb bacterial infection.

## Results

We demonstrate that total serum IgG N-glycosylation of acutely infected Lyme disease patients contrasts with the typical pro-inflammatory signature often found on other inflammatory diseases. First, a proof-of-concept study was performed using pooled serum to identify the IgG N-glycan signature of healthy control, acute LD, and patient-matched antibiotic-treated LD serum using high-pressure liquid chromatography separation (HPLC) and detection of fluorescently labeled glycans [69]. The samples were subsequently analyzed using the recently developed GlycoTyper MALDI method [70] which pairs a specific total-IgG capture antibody with subsequent MALDI-FT-ICR imaging (MALDI). The trends in the glycan signatures were reproducible between both platforms, and thus we proceeded to analyze a larger confirmatory set of individual serum samples using the GlycoTyper platform.

## Acute LD IgG N-glycans gain terminal galactose and sialic acid

HPLC analysis of the glycan signature revealed several statistically significant differences between control and acutely infected patients (**Fig 1**). Individual glycans were quantitated as a percent of the total glycan profile and compared across cohorts using one-way ANOVA (**Fig 1A**). Two agalactosylated glycan species, F(6)A2G0 and F(6)A2BG0, decreased in the acute and treated pooled cohorts when compared to controls. There was also an observed decrease in the mono-galactosylated F(6)A2G1 glycan. In addition, we observed significant increases in three glycan species. N-glycans containing terminal di-galactose, with and without core fucose increased significantly (A2G2, F(6)A2G2), as did the core fucosylated mono-sialylated glycan (F(6)A2G2S1). When the N-glycans were grouped by class (**Fig 1B**), the decrease in the smaller agalactosylated structures and an increase in the terminally galactosylated and sialic acid glycans continued to be observed. However, bisecting and core-fucosylated N-glycan classes did not significantly vary across cohorts. The only statistical difference observed between the acute and treated patient cohorts 90-day post-treatment was a continued increase in the addition of terminal galactose.

**Fig 1.**
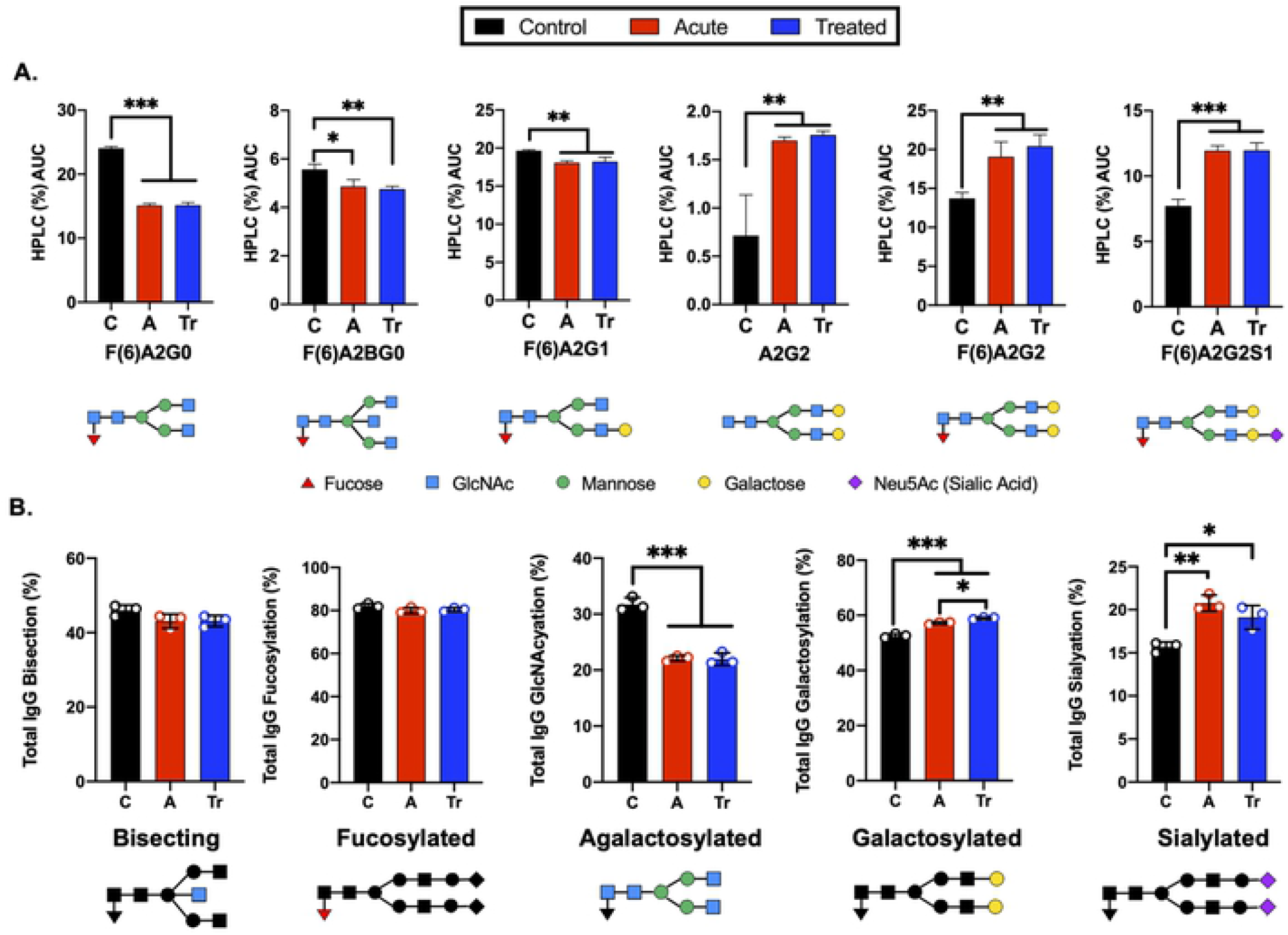
Analysis of IgG reveals significant changes in N-glycan structure distribution during acute and treated LD. **A)** Significant alterations of IgG N-glycans reported by HPLC analysis from pooled serum from control (C), acute (A), or treated (Tr) patients reported as the percent area under the curve (AUC) for each peak determined from the average of triplicate samples. N-glycan structures are displayed below. Control represents pooled healthy age-matched serum n=7, Acute represents pooled serum from two-tiered diagnosed Lyme disease patients n=5, Treated represents patients donating serum a second time 70-90 days after completion of the reportedly curative round of antibiotic treatment (14-21d doxycycline) for Lyme disease n=3. **B)** Labeled N-glycan classes: Bisecting, Fucosylated, Agalactosylated, Galactosylated, Sialylated detected using HPLC analysis of IgG N-glycans from 3 replicates +/- S.D. Analysis was completed using One-Way ANOVA with post-hoc Tukey’s multiple comparisons, *p<0.05, **p<0.01, ***p<0.001.

### MALDI-FT-ICR and HPLC detect similar trends of IgG N-glycans in LD

IgG N-glycans were desialylated and the HPLC analysis was compared to the MALDI-FT-ICR method. The IgG N-glycans were grouped by terminal sugar moiety into 5 glycan classes. These groups showed the same trends using both platforms (**Fig 2**). Significant reductions in agalactosylated N-glycans and significant increases in terminal galactose were identified in the HPLC (**Fig 2A**) and MALDI glycan analysis platforms (**Fig 2B**). Both methods indicate there is no significant difference in the abundance of bisecting or core-fucosylated N-glycans when comparing controls, acute and treated patient cohorts. The reproducible shifts in IgG N-glycan abundance within the desialylated glycan classes promoted the use of a larger confirmatory set of LD serum to be analyzed in a high-throughput manner using the MALDI method.

**Fig 2.**
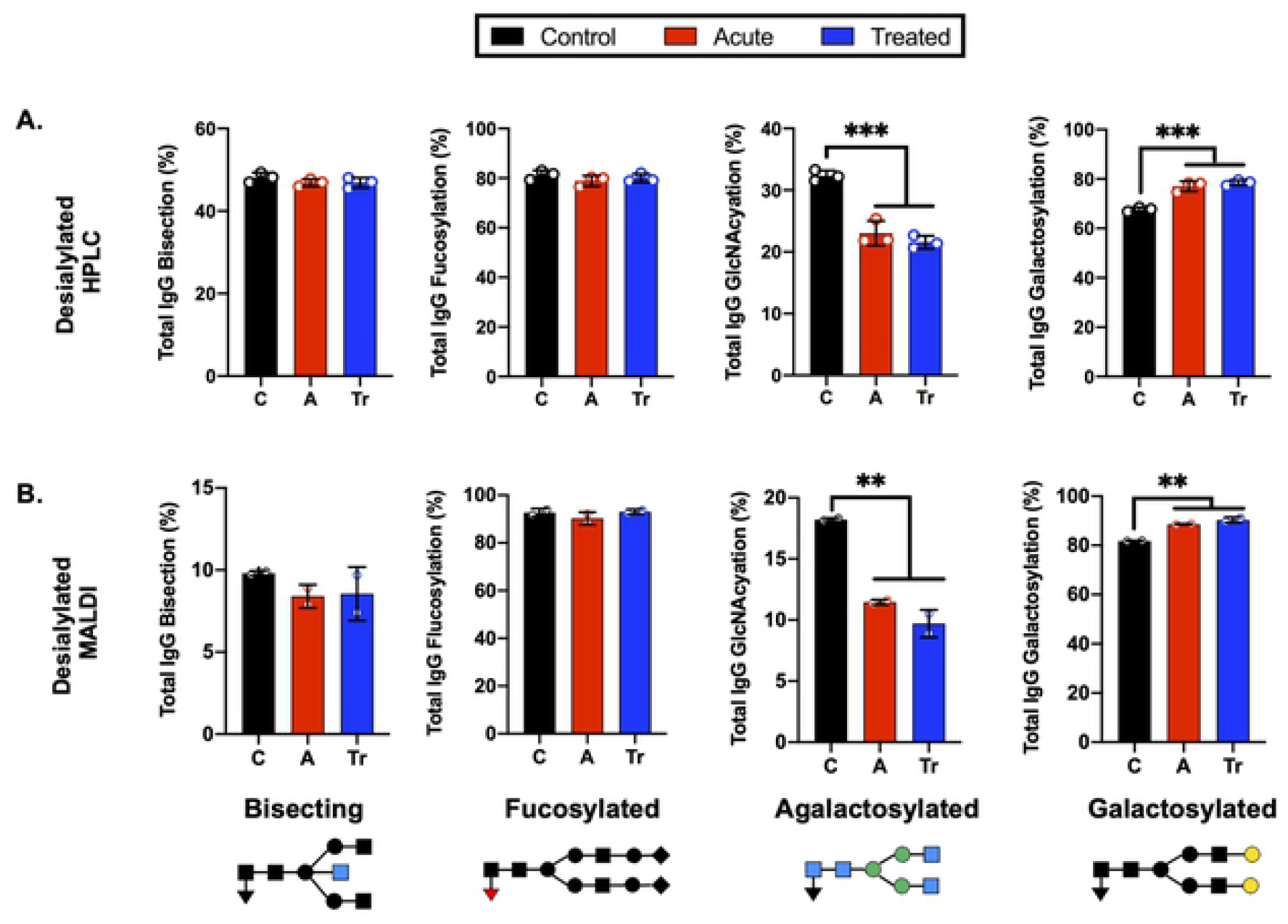
Grouping IgG N-glycans by class reveals MALDI and HPLC detect similar trends during LD. **A)** Labeled N-glycan classes: Bisecting, Fucosylated, Agalactosylated, Galactosylated, Sialylated detected using HPLC analysis of desialylated IgG N-glycans grouped by class averaged from 3 replicates of pooled cohorts described in Fig 1. **B)** MALDI analysis of desialylated IgG N-glycan classes averaged from 2 replicates of pooled cohorts described in Fig 1. Analysis was completed using One-Way ANOVA with post-hoc Tukey’s multiple comparisons, *p<0.05, **p<0.01, ***p<0.001.

### MALDI analysis identifies IgG N-glycosylation signatures of patients with LD

IgG N-glycans from a confirmatory set of healthy control, acute LD, and acute re-drawn serum post-antibiotic treatment patient sera were examined. MALDI analysis of the desialylated IgG N-glycans identified statistically significant shifts of N-glycan species (**Fig 4**). Several significant differences were detected when comparing control and acute LD.Acute LD patients exhibit a significant decrease in the core fucosylated agalactosylated N-glycan F(6)A2G0 when compared to controls (**Fig 3A**). Conversely, there was a statistically significant increase in the core fucosylated di-galactosylated N-glycan F(6)A2G2. IgG N-glycans terminating in total galactose or digalactose significantly increased during acute LD compared to healthy controls (**Fig 3B**). There was no significant difference in the total bisecting or core-fucosylated N-glycans when comparing Acute to healthy control IgG.

**Fig 3.**
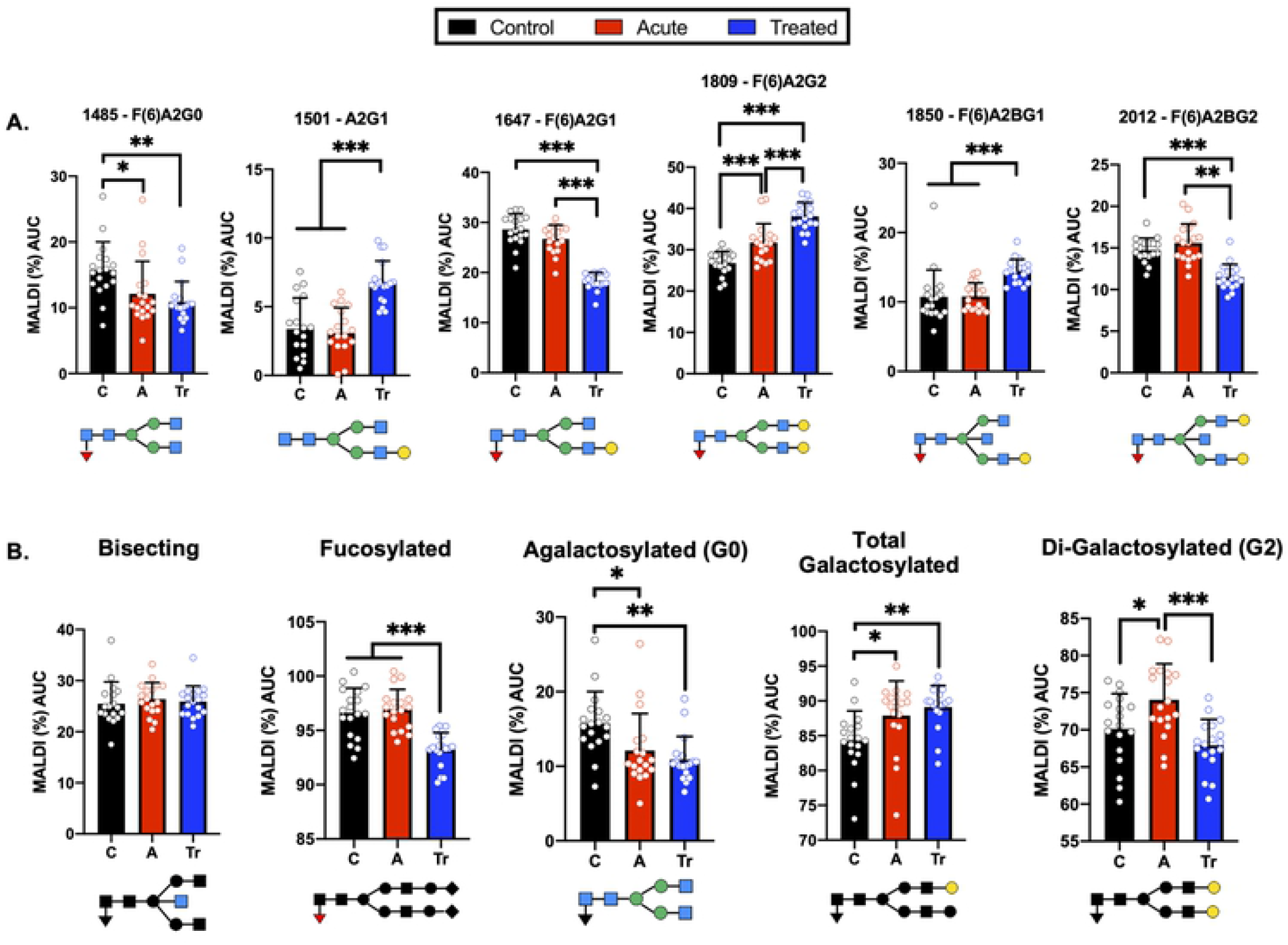
Confirmatory set of LD patient serum assayed using the high-throughput MALDI method. **A)** IgG N-glycans detected by MALDI analysis of Ab-captured IgG from n=18 cohorts individually run in triplicate +/- S.D. and reported as the percent of the total identified N-glycan m/z peak intensity. N-glycan structures displayed below **B)** MALDI analysis of desialylated IgG N-glycans grouped by class from n=18 samples per cohort averaged in triplicate +/- S.D. Labeled N-glycan class structures: Bisecting, Fucosylated, Agalactosylated, Galactosylated, and Di- Galactosylated are presented below the respective graph. Analysis was completed using One-Way ANOVA with post-hoc Tukey’s multiple comparisons, *p<0.05, **p<0.01, ***p<0.001. contain uncharacteristic increases in terminally galactosylated N-glycans.

We observed a continued perturbation of IgG glycosylation in the post-treatment cohort that does not return towards healthy control baselines for many detected N-glycans. Treated LD patients maintained a significant decrease in the agalactosylated F(6)A2G0 observed in acute LD (**Fig 3A**). An additional significant drop was seen in two glycan structures, F(6)A2G1 and F(6)A2BG2, when compared with control and acute. Several glycans showed significant increases. Increases that were limited to the treated cohort include the A2G1 and F(6)A2BG1 structures. The F(6)A2G2 continued to increase above the already elevated acute LD cohort level. Grouping the treated timepoint IgG N-glycans revealed a drop in total core-fucose as well as a decrease in the digalactosylated grouped N-glycans (**Fig 3B**). There was no difference observed for the total bisecting N-glycans between acute and treated timepoints.

These findings align with previous assays from the pooled pilot study experiments. Comparisons of IgG N-glycosylation dependence on age, sex, location of collection, or ethnicity were assessed. There was no statistically significant difference within or between cohorts when applying these metrics (data not shown) with one exception. Healthy control males had 2.3% higher F(6)A2G0 N-glycan profiles compared to their healthy control female counterparts. While IgG N-glycan signatures change during ageing and between sexes, the age- and sex-matched cohorts permitted comparisons across and between the cohorts [71].

### IgG N-glycans discriminate between healthy control and LD timepoints

Seroconversion is often employed as a LD diagnostic, yet there is great need for a more sensitive, early indicator of disease. Moreover, serological assays for LD cannot differentiate between a patient with acute LD compared to a patient that has recently convalesced from LD. **Fig 4** reports the efficacy of selected IgG N-glycans to discriminate between healthy and LD patient cohorts. Thresholds for listed N-glycan classes were determined using receiver operating characteristic (ROC) curve. The performance of the discrimination was reported in a confusion matrix (**Supplemental Fig 2-4**). Healthy controls are differentiated from acute serum samples using four N-glycan classes: F(6)A2G0, F(6)A2G2, percent total terminal galactose, and percent terminal di-galactose; resulting in 75% sensitivity, 100% specificity, and 85.7% accuracy. Healthy controls are differentiated from treated LD patient serum using the N-glycan classes: F(6)A2G0, A2G1, F(6)A2G1, F(6)A2G2, F(6)A2BG2, percent total fucose, and percent total terminal galactose; resulting in 100% sensitivity, 94.7% specificity, and 97.3% accuracy. Lastly, acute LD serum is differentiated from the treated serum using the N-glycan classes: A2G1, F(6)A2G1, F(6)A2G2, F(6)A2BG1, F(6)A2BG2, total percent fucose, and percent G2 galactosylation; resulting in 100% sensitivity, specificity, and accuracy.

**Fig 4.**
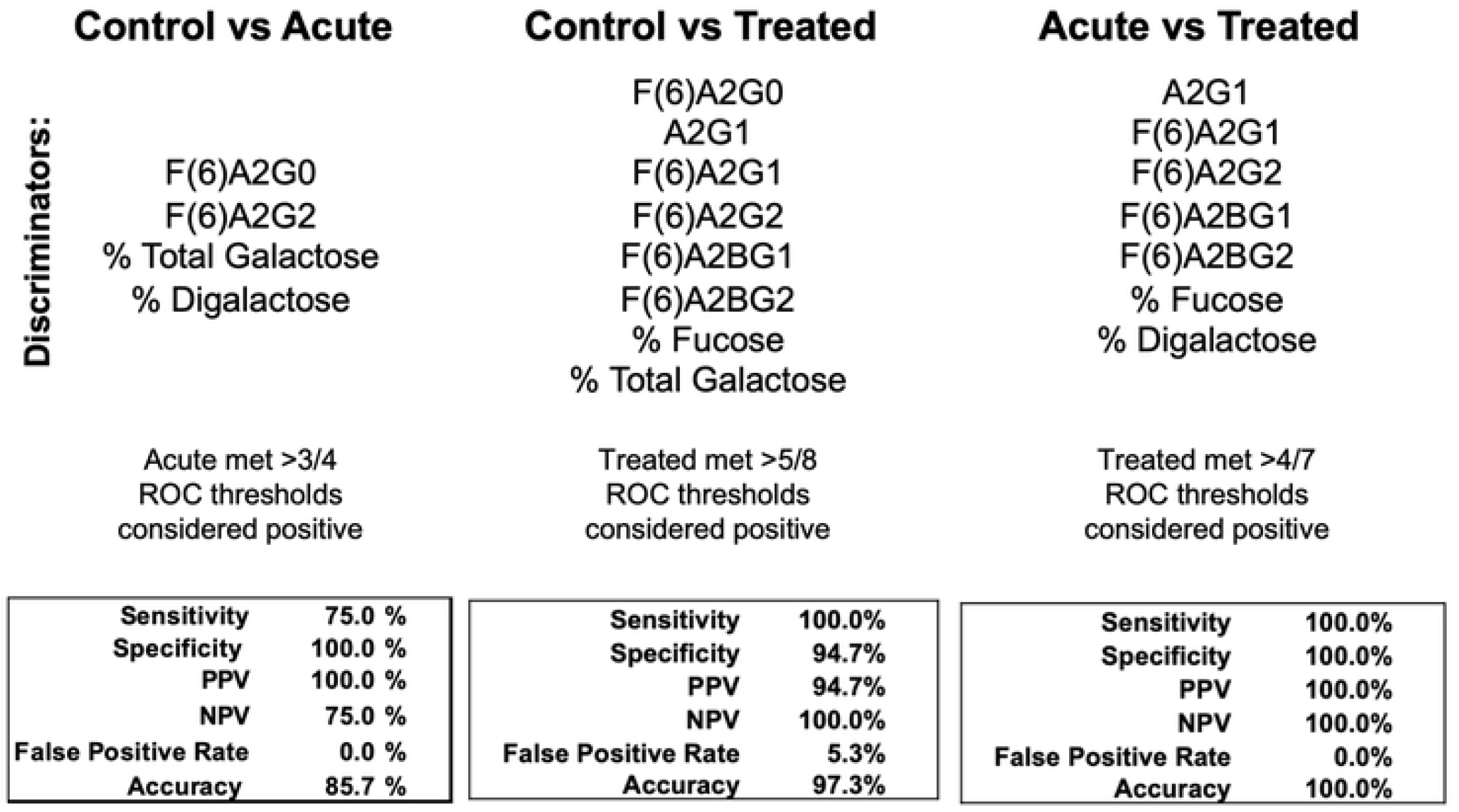
Confusion matrices from combined discriminatory IgG N-glycan ROC thresholds. N-glycans with significant discrimination between cohorts determined using ROC analysis are listed above. The listed N-glycan discriminators were combined for each cohort comparison (Control vs Acute LD, Control vs Treated LD, Treated vs Acute LD) and the resulting confusion matrices are reported.

### Lyme disease subverts IgG response – working hypothesis

Results presented in **Fig 5** show the increased terminal galactose observed on IgG N-glycans detected during LD within the context of the humoral immune response. IgG N-glycans detected in acute LD patients contain significantly higher levels of galactose and conversely lower agalactosylated structures. Most IgG N-glycans responding to inflammatory diseases present increased agalactosylated N-glycans to promote downstream pro-inflammatory immune responses. Thus, the LD IgG N-glycans may be induced through the Borrelia *burgdorferi’s* disruption of the germinal center and antibody response.

**Fig 5.**
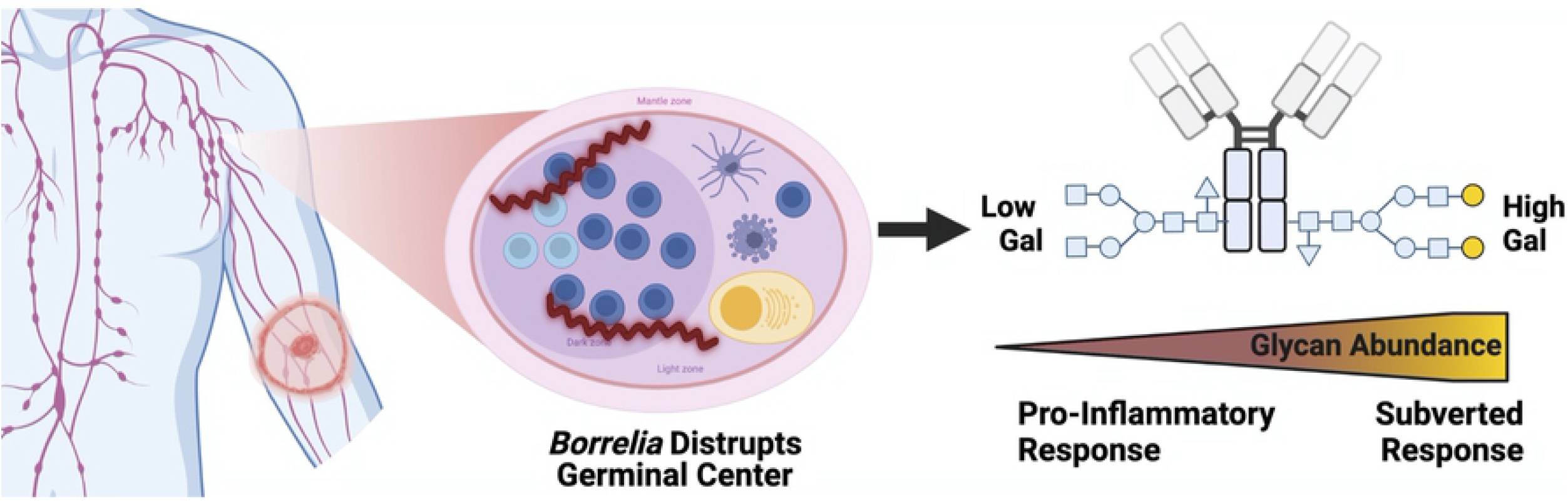
IgG N-glycans Respond to acute Lyme disease infection with increased galactose, impairing bound IgG to signal for inflammation, unlike other pro-inflammatory disease IgG signatures. Left to right, a tick infected with *Borrelia* transmits the infection to the human host leading to LD. The Bb spirochete disrupts the architecture of the germinal center within the lymph node. IgG molecules produced.

## Discussion

This is the first report of total human IgG N-glycan profiles during early and treated LD. Our initial studies detected an unanticipated shift in IgG N-glycan profile which was confirmed and expanded upon using a larger set of samples. During many inflammatory states (**Table 1**), IgG N-glycan profiles respond with reduced terminal sialic acids and galactose and increased GlcNAc sugar exposure [40]. This agalactosylated IgG phenotype has been demonstrated to promote pro-inflammatory responses and may aid in clearing the infection [44–50, 72, 73]. In the case of LD, total IgG N-glycan profiles had the opposite trend. LD increases the galactose and sialic acid content of IgG N-glycans while further decreasing the terminal GlcNAc exposure (**Fig 1–4**). The result of such a shift requires further investigation, but in theory could contribute to ineffective host responses to LD infection by reducing humoral immune activation [72, 73]. The ability to detect these changes within total IgG suggests that LD induces a large shift in the IgG N-glycan composition. Thus, this dysregulated immune response may at least in part explain why human IgG produced during an initial LD infection is not effective to clear the current infection or protect from a future re-infection with LD.

The pilot study served to compare the well-established HPLC IgG N-glycan analysis method [74] to the novel MALDI analysis method [75] using pooled cohorts of control, acute LD, and treated LD serum. Due to MALDI’s limited detection of unmodified sialic acids, samples were treated with a sialidase [76]. The results presented in **Fig 2** demonstrate that both methods detect comparable individual identified N-glycans with similar trends of terminal galactose exposure induced during LD and maintained during the post-antibiotic treatment time points. To confirm the pooled pilot study findings, three cohorts of 18 individual samples were analyzed in triplicate using the MALDI method. We found that the demographics of the age- and sex-matched patient cohorts did not lead to inherent differences in IgG N-glycosylation.

While the MALDI results confirmed the original trends observed in the pooled pilot study, a higher number of samples analyzed led to further trends emerging (**Fig 3A, 3B)**. Agalactosylated structures are reduced while total N-glycans terminating with a galactose moiety continued to increase within the acute and treated cohorts as indicated within the pooled pilot experiments. N-glycans containing a core-fucose are markedly lowered in the antibiotic-treated cohort which suggests an increase in Antibody-Dependent Cellular Cytotoxicity (ADCC) abilities of post-treatment IgG [77]. The increased level of digalactosylated (G2) N-glycans on IgG at the acute stage and subsequent return to healthy control levels is a potential biomarker reflecting the host’s response to successful antibiotic treatment. Following antibiotic therapy for Tuberculosis (TB), IgG N-glycan diagalactose content returned to healthy control levels [51]. Grace et al. found G2 N-glycans decreased during acute TB and subsequently increased after effective antibiotic treatment. In the case of LD, IgG N-glycans increase G2 content during infection and return to healthy control levels after effective antibiotic treatment (**Fig 3B)**. This once more converse trend observed in LD suggests a subverted immune response during LD compared to TB infection. The implication of the IgG N-glycan response to LD is portrayed in **Fig 5**.

LD patients are discriminated from healthy controls with a high degree of sensitivity (75-100%) and specificity (94.7-100%) using total IgG N-glycan measurements (**Fig 4**). Blinded analysis of acute LD, post-treatment LD, and mimic diseases using these methods should be completed to validate these findings. Future LD tests incorporating total IgG N-glycan analysis could increase acute LD diagnostic sensitivity and track subsequent antibiotic treatment responses.

Mechanisms controlling the dynamic B-cell glycosyltransferase expression operating on IgG N-glycans during LD require examination. B-cell glycosyltransferase expression is known to respond to the cytokine micro-environment [78]. Recent studies detected IgG N-glycosylation profiles are impacted by the type of adjuvant present during vaccination [79]. Additionally, during the onset of autoimmune disease, T_H_17 cells signal newly differentiated B cells with IL-22 and IL- 23 to regulate glycosyltransferase expression, resulting in a pro-inflammatory agalactosylated, non-fucosylated IgG N-glycan [80]. The increase in terminal galactose of IgG N-glycans during acute LD may be attributed to an upregulation of naïve B-cell beta-1,4-galactosyltransferase expression [68, 81–83]. Future studies should determine if LD induces a specific cytokine signaling pathway to affect the glycosyltransferase expression of plasma cells or B cells during LD infection.

Lyme disease has been demonstrated to destroy germinal centers of lymph nodes during early infection [72, 73, 84, 85]. These germinal centers are a vital structure that produces long-lived immunoglobulin responses through T-cell-dependent interactions [86]. The destruction of the germinal center may explain why patients are liable to become re-infected with LD after treatment. Additionally, Bb has been demonstrated to gain entry inside human endothelial cells [87], alter innate immune responses after phagocytosis [88], suppress lymphocytes growth rate [89], and down-regulate major-histocompatibility complexes expression on Langerhans cells [90]. Any one of these effects could be responsible for altering the IgG N-glycan profile during LD. For example, antigens presented in a T-cell *independent* manner promote increased sialyation on the Fc portion of IgG leading to an immunotolerant, less pro-inflammatory IgG N-glycan repertoire [91]. Interestingly, murine studies have demonstrated a reduction in the general humoral response after LD infection as indicated by low titers of anti-viral antibodies produced post-vaccination and Bb-impaired helper T-cell mediated affinity maturation [84, 92]. Lastly, because the LD cohort completed 2-3 week of oral doxycycline therapy, the serum N-glycome or immune response could have been altered in part due to changes in gut flora [93, 94].

## Conclusion

Using the GlycoTyper MALDI-FT-ICR imaging approach, we detected an unexpected IgG N-glycan signature in humans during LD. We have demonstrated the IgG N-glycans produced during a Lyme disease infection lack the classic highly agalactosylated signature associated with most inflammatory diseases. Instead, LD induces IgG N-glycans with larger, terminally galactosylated sugar moieties. Moreover, many IgG N-glycans detected at the acute LD timepoint remain elevated or altered at the post-antibiotic treatment timepoint. Fascinatingly, the N-glycans terminating in digalactose returned to a healthy baseline after antibiotic treatment. Furthermore, we detected a significant decrease in total core-fucosylation at the treated timepoint suggesting a possible increase in ADCC which may aid in clearing Bb from the host. IgG N-glycans offer numerous biomarkers, reflect acute disease state, response to treatment, and may improve the sensitivity of the acute diagnosis of LD above the current two-tiered testing protocol. This first examination of IgG N-glycan signatures associated with LD requires future study.

## Material and methods

### Patient samples

Serum samples (**Table 2**) were obtained from the Bay Area Lyme Disease Biobank and stored at −80°C. Serology was determined at Stony Brook University [95] and the Bay Area Lyme Disease Biobank. Human subject research IRB requirements were met (IRB #1808006553). All LD patients presented with erythema migrans (EM) rash(es) at the time of diagnosis and were confirmed positive for LD using the two-tiered serological studies or PCR identification. Convalescent draws were obtained after patients completed their prescribed course of 14-21 days of antibiotics and convalesced for 70-90 days without further symptoms. In the pilot study, serum was pooled into three patient cohorts: healthy control (n=7), acute LD (n=5), and treated LD serum (n=3). In the subsequent GlycoTyper study, data was collected from each individual patient: healthy control (n=18), acute (n=18), and patient-matched treated (n=18). Further demographic details for the patient cohorts are presented in **Supplemental Table 1**.

**Table 2:**
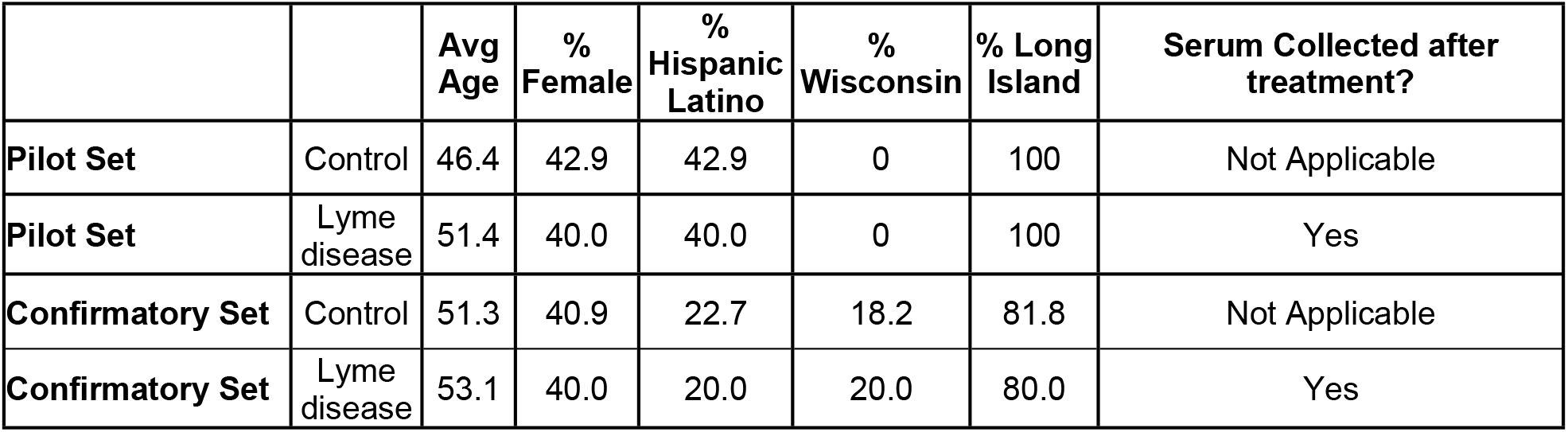
Patient samples. Table 2 legend. Healthy Control and Lyme disease LD with matched Acute and Post-treatment Demographics of Bay Area Lyme Disease Biobank Serum Samples. Sample ID numbers are linked to demographic details including age, sex, ethnicity, antibiotics (Abx) prescribed, and if the patient donated a post-treatment convalescent serum sample. Samples assayed within the pilot study and/or the later N=18 study are denoted.

### HPLC IgG N-glycan analysis

IgG was purified from 5μL of pooled serum using Protein A/G UltraLink Resin (Thermo Scientific, MA) according to the manufacturer’s directions. IgG Heavy Chains (Fc region) were isolated using 1D gel electrophoresis, stained with Coomassie stain and the 50kDa band was excised. Following detaining, the glycans were enzymatically removed and fluorescently labeled following standard in-gel PNGase F and labeling protocols as previously described [96, 97]. The labeled N-glycans were combined with 100% Acetonitrile (30:70) in an HPLC-compatible vial. Fluorescently labeled glycans were subsequently analyzed by high-performance liquid chromatography (HPLC) by using a TSK-Gel Amide 80 column (Tosoh Bioscience LLC). The mobile phase consisted of solvent A (50 mmol/L ammonium formate, pH 4.4) and solvent B (acetonitrile). The gradient used was as follows: a linear gradient from 20% to 58% solvent A at 0.4 mL/min for 152 min followed by a linear gradient from 58% to 100% solvent A for the next 3 minutes. The flow rate was increased to 1.0 mL/min, and the column was washed in 100% solvent A for 5 minutes. Following the wash step, the column was equilibrated in 20% solvent A for 22 minutes in preparation for the next sample run. HPLC analysis was performed using the Waters Alliance HPLC System, complemented with a Waters fluorescence detector, and quantified by the Millennium Chromatography Manager (Waters Corporation). Glycan structures were identified by calculating the glucose unit and GlycoStore database as previously described [98] HPLC method overview is provided in **Supplemental Fig 1A**.

### Removal of sialic acids for HPLC and MALDI comparison

Due to the inherent challenges of sialic acid containing glycans in mass spectrometry, HPLC data was collected using samples that were desialylated to allow a direct comparison with the MALDI-FT-ICR method. Beginning with 13μL of the labeled and chromatographically cleaned 2-AB glycans, 4μL 5X pH 6.0 Enzyme Buffer was added before pipette mixing 3μL of Sialidase A (ProZyme (now AdvanceBio), OH). The 20μL final volume was incubated at 32°C for 12 hours. Next, the 20μL solution was added to a 10K MWCO concentrator column and centrifuged at 12,000 rpm for 10 minutes. The flow-through was collected and combined with an additional 25μL of dH2O before being added back to the 10K MWCO column and centrifuged at 12,000 rpm. This process of collecting the flow-through and combining it with additions of 25μL dH2O was repeated to serially enrich the labeled N-glycans while filtering out the sialidase enzyme. The sample was subsequently dried down via SpeedVac holding a vacuum at −28in Hg without added heat and re-suspended in 30μL dH2O. Sialidase-treated samples were then analyzed as outlined for non-sialidase-treated sample in HPLC.

### MALDI-FT-ICR mass spectrometry N-glycan analysis

SolarisX Legacy 7T FT-ICR mass spectrometer equipped with a Matrix-Assisted Laser Desorption/Ionization (MALDI) (Bruker) analysis of total IgG N-glycans is detailed in the literature [56, 70, 76, 99]. In brief, serum samples incubated with 0.2μg/μl spotted anti-IgG capture antibodies (Bethyl Laboratories Inc., Tx, Cat. Number A80-104) treated with Sialidase A (produced in-house by MUSC), sprayed with PNGase F (produced by N-Zyme, PA) to release N-glycans from captured targets, coated with a matrix, and analyzed for glycan abundance at specific m/z peaks by MALDI-FT-ICR MS using SCiLS Lab software 2022a (Bruker). A capture antibody treated with PBS served as a blank to subtract the N-glycans released from the capture antibody from the final analysis. A MALDI-FT-ICR (referred to as MALDI) method overview is provided in **Supplemental Fig 1B.**

## Statistical analysis

One-way ANOVA analysis with post-hoc Tukey’s multiple comparisons test was employed to examine the triplicate datasets for statistically significant differences between cohorts. P<0.05 was considered statistically significant and figures are denoted as having *p<0.05, **p<0.01, ***p<0.001. All figures and analyzes were completed using GraphPad Prism 8. Results are described using the Oxford nomenclature. [100]

## Acknowledgments

We thank the following individuals: Dr. Laura Steel and Mr. Mengjun Wang for their support. We also would like to acknowledge Dr. Elizabeth Horn, Bay Area Lyme Disease Biobank.

## Supporting information

**Supplementary Fig 1. Method comparison between HPLC and MALDI. (A)** HPLC methodology to identify and quantitate Protein A/G purified IgG N-glycan released from patient cohort pooled serum. **(B)** MALDI MS method to identify and quantitate Ab-captured IgG N-glycans released from patient cohort pooled serum.

**Supplemental Fig 2. Acute versus Control Selected N-glycan class discriminators.** Four N-glycan classes: F(6)A2G0, F(6)A2G2, % Total Galactose, and % Terminal Digalactose were used in combination to discriminate between the healthy age- and sex-matched healthy controls compared to the Acute LD patients. A confusion matrix was tabulated from the performance of the 3 out of 4 positive (value highlighted in pink) criteria.

**Supplemental Fig 3. Acute versus Treated Selected N-glycan class discriminators.** Seven N-glycan classes were used in combination to discriminate between the Acute LD compared to the Treated LD timepoints. A confusion matrix was tabulated from the performance of the 4 out of 7 positive (value highlighted in pink) criteria.

**Supplemental Fig 4. Treated versus Control Selected N-glycan class discriminators.** Two N-glycan classes were used in combination to discriminate between the age- and sex-matched healthy controls compared to the Treated LD cohort. A confusion matrix was tabulated from the performance of the 2 out of 2 positive (value highlighted in pink) criteria.

**Supplemental Table 1. Demographics of Bay Area Lyme Disease Biobank Serum Samples.** Supplementary table legend: Sample ID numbers are linked to demographic details including age, sex, ethnicity, antibiotics (Abx) prescribed, and if the patient donated a post-treatment convalescent serum sample. Samples assayed within the pilot study and/or the later N=18 study are noted.

## Notes

### Competing Interest Statement

Anand Mehta is a founding partner in GlycoPath Inc. and holds patents for the GlycoTyper methodology.

## References

1. Kugeler, K.J., et al., Estimating the Frequency of Lyme Disease Diagnoses, United States, 2010–2018. Emerging Infectious Diseases, 2021. 27(2): p. 616–619.

2. Hugli, D., et al., Tick bites in a Lyme borreliosis highly endemic area in Switzerland. International Journal of Medical Microbiology, 2009. 299(2): p. 155–160.

3. Dhruv and Charles, Spatiotemporal Evolution of Erythema Migrans, the Hallmark Rash of Lyme Disease. Biophysical Journal, 2014. 106(3): p. 763–768.

4. Donta, S.T., et al., Report of the Pathogenesis and Pathophysiology of Lyme Disease Subcommittee of the HHS Tick Borne Disease Working Group. Front Med (Lausanne), 2021. 8: p. 643235.

5. Bender, P.D. and J.S. Ilgen, Early disseminated Lyme disease. BMJ Case Reports, 2018: p. bcr-2017-223889.

6. Mac, S., S.R. Da Silva, and B. Sander, The economic burden of Lyme disease and the cost-effectiveness of Lyme disease interventions: A scoping review. PLOS ONE, 2019. 14(1): p. e0210280.

7. Zhang, X., et al., Economic Impact of Lyme Disease. Emerging Infectious Diseases, 2006. 12(4): p. 653–660.

8. Davidsson, M., The Financial Implications of a Well-Hidden and Ignored Chronic Lyme Disease Pandemic. Healthcare (Basel), 2018. 6(1).

9. Maloney, E.L., Evidence-Based, Patient-Centered Treatment of Erythema Migrans in the United States. Antibiotics, 2021. 10(7): p. 754.

10. Cameron, D.J., L.B. Johnson, and E.L. Maloney, Evidence assessments and guideline recommendations in Lyme disease: the clinical management of known tick bites, erythema migrans rashes and persistent disease. Expert Review of Anti-infective Therapy, 2014. 12(9): p. 1103–1135.

11. Roome, A., et al., Lyme Disease Transmission Risk: Seasonal Variation in the Built Environment. Healthcare, 2018. 6(3): p. 84.

12. Bhatia, B., et al., Infection history of the blood-meal host dictates pathogenic potential of the Lyme disease spirochete within the feeding tick vector. PLOS Pathogens, 2018. 14(4): p. e1006959.

13. Norman, M.U., et al., Molecular Mechanisms Involved in Vascular Interactions of the Lyme Disease Pathogen in a Living Host. PLoS Pathogens, 2008. 4(10): p. e1000169.

14. Casselli, T., et al., A murine model of Lyme disease demonstrates that Borrelia burgdorferi colonizes the dura mater and induces inflammation in the central nervous system. PLOS Pathogens, 2021. 17(2): p. e1009256.

15. Halperin, J.J., Nervous System Lyme Disease: Diagnosis and Treatment. Current Treatment Options in Neurology, 2013. 15(4): p. 454–464.

16. Lelovas, P., et al., Cardiac implications of Lyme disease, diagnosis and therapeutic approach. Int J Cardiol, 2008. 129(1): p. 15–21.

17. Arvikar, S.L. and A.C. Steere, Diagnosis and Treatment of Lyme Arthritis. Infectious Disease Clinics of North America, 2015. 29(2): p. 269–280.

18. Berglund, J., et al., An Epidemiologic Study of Lyme Disease in Southern Sweden. New England Journal of Medicine, 1995. 333(20): p. 1319–1324.

19. Cardenas-de la Garza, J.A., et al., Clinical spectrum of Lyme disease. Eur J Clin Microbiol Infect Dis, 2019. 38(2): p. 201–208.

20. Bratton, R.L., et al., Diagnosis and treatment of Lyme disease. Mayo Clin Proc, 2008. 83(5): p. 566–71.

21. Shor, S., et al., Chronic Lyme Disease: An Evidence-Based Definition by the ILADS Working Group. Antibiotics, 2019. 8(4): p. 269.

22. Maloney, E.L., Controversies in Persistent (Chronic) Lyme Disease. Journal of Infusion Nursing, 2016. 39(6): p. 369–375.

23. Marques, A.R., Laboratory Diagnosis of Lyme Disease. Infectious Disease Clinics of North America, 2015. 29(2): p. 295–307.

24. Hyde, J.A., Borrelia burgdorferi Keeps Moving and Carries on: A Review of Borrelial Dissemination and Invasion. Front Immunol, 2017. 8: p. 114.

25. Lantos, P.M., P.G. Auwaerter, and C.A. Nelson, Lyme Disease Serology. JAMA, 2016. 315(16): p. 1780.

26. Tokarz, R., et al., A multiplex serologic platform for diagnosis of tick-borne diseases. Sci Rep, 2018. 8(1): p. 3158.

27. Chou, E., et al., A fluorescent plasmonic biochip assay for multiplex screening of diagnostic serum antibody targets in human Lyme disease. PLoS One, 2020. 15(2): p. e0228772.

28. Schutzer, S.E., et al., Direct Diagnostic Tests for Lyme Disease. Clinical Infectious Diseases, 2019. 68(6): p. 1052–1057.

29. Garg, K., et al., Evaluating polymicrobial immune responses in patients suffering from tick-borne diseases. Sci Rep, 2018. 8(1): p. 15932.

30. Moore, A., et al., Current Guidelines, Common Clinical Pitfalls, and Future Directions for Laboratory Diagnosis of Lyme Disease, United States. Emerg Infect Dis, 2016. 22(7).

31. Bobe, J.R., et al., Recent Progress in Lyme Disease and Remaining Challenges. Front Med (Lausanne), 2021. 8: p. 666554.

32. Parekh, R., et al., A comparative analysis of disease-associated changes in the galactosylation of serum IgG. Journal of Autoimmunity, 1989. 2(2): p. 101–114.

33. Parekh, R.B., et al., Association of rheumatoid arthritis and primary osteoarthritis with changes in the glycosylation pattern of total serum IgG. Nature, 1985. 316(6027): p. 452–457.

34. Sun, D., et al., Distribution of abnormal IgG glycosylation patterns from rheumatoid arthritis and osteoarthritis patients by MALDI-TOF-MSn. The Analyst, 2019. 144(6): p. 2042–2051.

35. Trbojević Akmačić, I., et al., Inflammatory Bowel Disease Associates with Proinflammatory Potential of the Immunoglobulin G Glycome. Inflammatory Bowel Diseases, 2015: p. 1.

36. Clerc, F., et al., Human plasma protein N-glycosylation. Glycoconjugate Journal, 2016. 33(3): p. 309–343.

37. Varki, A., Biological roles of glycans. Glycobiology, 2017. 27(1): p. 3–49.

38. Varki, A., Sialic acids in human health and disease. Trends in Molecular Medicine, 2008. 14(8): p. 351–360.

39. Cobb, B.A., The history of IgG glycosylation and where we are now. Glycobiology, 2020. 30(4): p. 202–213.

40. Gudelj, I., G. Lauc, and M. Pezer, Immunoglobulin G glycosylation in aging and diseases. Cellular Immunology, 2018. 333: p. 65–79.

41. Mahan, A.E., et al., A method for high-throughput, sensitive analysis of IgG Fc and Fab glycosylation by capillary electrophoresis. J Immunol Methods, 2015. 417: p. 34–44.

42. Baum, L.G. and B.A. Cobb, The direct and indirect effects of glycans on immune function. Glycobiology, 2017. 27(7): p. 619–624.

43. Hitsumoto, Y., et al., Relationship Between Interleukin 6, Agalactosyl Igg And Pristane-Induced Arthritis. Autoimmunity, 1992. 11(4): p. 247–254.

44. Dekkers, G., T. Rispens, and G. Vidarsson, Novel Concepts of Altered Immunoglobulin G Galactosylation in Autoimmune Diseases. Front Immunol, 2018. 9: p. 553.

45. Dekkers, G., et al., Decoding the Human Immunoglobulin G-Glycan Repertoire Reveals a Spectrum of Fc-Receptor-and Complement-Mediated-Effector Activities. Frontiers in Immunology, 2017. 8(877).

46. Biermann, M.H.C., et al., Sweet but dangerous – the role of immunoglobulin G glycosylation in autoimmunity and inflammation. Lupus, 2016. 25(8): p. 934–942.

47. Collin, M. and M. Ehlers, The carbohydrate switch between pathogenic and immunosuppressive antigen-specific antibodies. Experimental Dermatology, 2013. 22(8): p. 511–514.

48. De Jong, S.E., et al., IgG1 Fc N-glycan galactosylation as a biomarker for immune activation. Scientific Reports, 2016. 6(1): p. 28207.

49. Tijardović, M., et al., Intense Physical Exercise Induces an Anti-inflammatory Change in IgG N-Glycosylation Profile. Frontiers in Physiology, 2019. 10(1522).

50. Alter, G., T.H.M. Ottenhoff, and S.A. Joosten, Antibody glycosylation in inflammation, disease and vaccination. Semin Immunol, 2018. 39: p. 102–110.

51. Grace, P.S., et al., Antibody Subclass and Glycosylation Shift Following Effective TB Treatment. Front Immunol, 2021. 12: p. 679973.

52. Bond, A., et al., A Detailed Lectin Analysis of IgG Glycosylation, Demonstrating Disease Specific Changes in Terminal Galactose and N-acetylglucosamine. Journal of Autoimmunity, 1997. 10(1): p. 77–85.

53. Gardinassi Luiz, G., et al., Clinical Severity of Visceral Leishmaniasis Is Associated with Changes in Immunoglobulin G Fc N-Glycosylation. mBio. 5(6): p. e01844–14.

54. Ho, C.-H., et al., Aberrant Serum Immunoglobulin G Glycosylation in Chronic Hepatitis B Is Associated With Histological Liver Damage and Reversible by Antiviral Therapy. Journal of Infectious Diseases, 2015. 211(1): p. 115–124.

55. Mehta, A.S., et al., Increased Levels of Galactose-Deficient Anti-Gal Immunoglobulin G in the Sera of Hepatitis C Virus-Infected Individuals with Fibrosis and Cirrhosis. Journal of Virology, 2008. 82(3): p. 1259–1270.

56. Scott, D.A., et al., GlycoFibroTyper: A Novel Method for the Glycan Analysis of IgG and the Development of a Biomarker Signature of Liver Fibrosis. Frontiers in Immunology, 2022. 13.

57. Lin, S., et al., Serum immunoglobulin G N-glycome: a potential biomarker in endometrial cancer. Annals of Translational Medicine, 2020. 8(12): p. 748–748.

58. Vučković, F.’ et al., Association of Systemic Lupus Erythematosus With Decreased Immunosuppressive Potential of the IgG Glycome. Arthritis & Rheumatology, 2015. 67(11): p. 2978–2989.

59. Vicente, M.M., et al., Altered IgG glycosylation at COVID-19 diagnosis predicts disease severity. European Journal of Immunology, 2022.

60. Fokkink, W.J.R., et al., Immunoglobulin G Fc N-glycosylation in Guillain-Barré syndrome treated with intravenous immunoglobulin. Clinical & Experimental Immunology, 2014. 178: p. 105–107.

61. Blundell, P.A., et al., Engineering the fragment crystallizable (Fc) region of human IgG1 multimers and monomers to fine-tune interactions with sialic acid-dependent receptors. Journal of Biological Chemistry, 2017. 292(31): p. 12994–13007.

62. Pleass, R.J., The therapeutic potential of sialylated Fc domains of human IgG. mAbs, 2021. 13(1): p. 1953220.

63. Malphettes, L., et al., Highly efficient deletion of FUT8 in CHO cell lines using zinc-finger nucleases yields cells that produce completely nonfucosylated antibodies. Biotechnology and Bioengineering, 2010. 106(5): p. 774–783.

64. Beck, A. and J.M. Reichert, Marketing approval of mogamulizumab. mAbs, 2012. 4(4): p. 419–425.

65. Pereira, N.A., et al., The “less-is-more”in therapeutic antibodies: Afucosylated anti-cancer antibodies with enhanced antibody-dependent cellular cytotoxicity. mAbs, 2018. 10(5): p. 693–711.

66. Mimura, Y., et al., Glycosylation engineering of therapeutic IgG antibodies: challenges for the safety, functionality and efficacy. Protein & Cell, 2018. 9(1): p. 47–62.

67. Alter, G., T.H.M. Ottenhoff, and S.A. Joosten, Antibody glycosylation in inflammation, disease and vaccination. Seminars in Immunology, 2018. 39: p. 102–110.

68. Irvine, E.B. and G. Alter, Understanding the role of antibody glycosylation through the lens of severe viral and bacterial diseases. Glycobiology, 2020. 30(4): p. 241–253.

69. Campbell, M.P., et al., GlycoBase and autoGU: tools for HPLC-basedglycan analysis. Bioinformatics, 2008. 24(9): p. 1214–1216.

70. Black, A.P., et al., Antibody Panel Based N- Glycan Imaging for N- Glycoprotein Biomarker Discovery. Current Protocols in Protein Science, 2019. 98(1).

71. Yu, X., et al., Profiling IgG N-glycans as potential biomarker of chronological and biological ages: A community-based study in a Han Chinese population. Medicine (Baltimore), 2016. 95(28): p. e4112.

72. Hastey, C.J., et al., Delays and Diversions Mark the Development of B Cell Responses toBorrelia burgdorferiInfection. The Journal of Immunology, 2012. 188(11): p. 5612–5622.

73. Tunev, S.S., et al., Lymphoadenopathy during Lyme Borreliosis Is Caused by Spirochete Migration-Induced Specific B Cell Activation. PLoS Pathogens, 2011. 7(5): p. e1002066.

74. Comunale, M.A., et al., Total serum glycan analysis is superior to lectin-FLISA for the early detection of hepatocellular carcinoma. PROTEOMICS-Clinical Applications, 2013. 7(9-10): p. 690–700.

75. Black, A.P., et al., Antibody Panel Based N-Glycan Imaging for N-Glycoprotein Biomarker Discovery. Curr Protoc Protein Sci, 2019. 98(1): p. e99.

76. Black, A.P., et al., A Novel Mass Spectrometry Platform for Multiplexed N-Glycoprotein Biomarker Discovery from Patient Biofluids by Antibody Panel Based N-Glycan Imaging. Anal Chem, 2019. 91(13): p. 8429–8435.

77. Shields, R.L., et al., Lack of Fucose on Human IgG1 N-Linked Oligosaccharide Improves Binding to Human FcγRIII and Antibody-dependent Cellular Toxicity. Journal of Biological Chemistry, 2002. 277(30): p. 26733–26740.

78. Cao, Y., et al., Cytokines in the Immune Microenvironment Change the Glycosylation of IgG by Regulating Intracellular Glycosyltransferases. Frontiers in Immunology, 2022. 12.

79. Bartsch, Y.C., et al., IgG Fc sialylation is regulated during the germinal center reaction following immunization with different adjuvants. Journal of Allergy and Clinical Immunology, 2020. 146(3): p. 652–666.e11.

80. Pfeifle, R., et al., Regulation of autoantibody activity by the IL-23–TH17 axis determines the onset of autoimmune disease. Nature Immunology, 2017. 18(1): p. 104–113.

81. Keusch, J., P.M. Lydyard, and P.J. Delves, The effect on IgG glycosylation of altering 1,4-galactosyltransferase-1 activity in B cells. Glycobiology, 1998. 8(12): p. 1207–1213.

82. Dekkers, G., et al., Multi-level glyco-engineering techniques to generate IgG with defined Fc-glycans. Scientific Reports, 2016. 6(1): p. 36964.

83. Omtvedt, L.A., et al., Glycan analysis of monoclonal antibodies secreted in deposition disorders indicates that subsets of plasma cells differentially process IgG glycans. Arthritis & Rheumatism, 2006. 54(11): p. 3433–3440.

84. Elsner, R.A., et al., Suppression of Long-Lived Humoral Immunity Following Borrelia burgdorferi Infection. PLOS Pathogens, 2015. 11(7): p. e1004976.

85. Anderson, C. and C.A. Brissette, The Brilliance of Borrelia: Mechanisms of Host Immune Evasion by Lyme Disease-Causing Spirochetes. Pathogens, 2021. 10(3): p. 281.

86. Nothelfer, K., P.J. Sansonetti, and A. Phalipon, Pathogen manipulation of B cells: the best defence is a good offence. Nature Reviews Microbiology, 2015. 13(3): p. 173–184.

87. Ma, Y., A. Sturrock, and J.J. Weis, Intracellular localization of Borrelia burgdorferi within human endothelial cells. Infect Immun, 1991. 59(2): p. 671–8.

88. Petnicki-Ocwieja, T. and A. Kern, Mechanisms of Borrelia burgdorferi internalization and intracellular innate immune signaling. Front Cell Infect Microbiol, 2014. 4: p. 175.

89. Chiao, J.W., et al., Antigens of Lyme disease of spirochaeteBorrelia burgdorferiinhibits antigen or mitogen-induced lymphocyte proliferation. FEMS Immunology & Medical Microbiology, 1994. 8(2): p. 151–155.

90. Aberer, E., F. Koszik, and M. Silberer, Why Is Chronic Lyme Borreliosis Chronic? Clinical Infectious Diseases, 1997. 25(s1): p. S64–S70.

91. Oefner, C.M., et al., Tolerance induction with Tcell–dependentprotein antigens induces regulatory sialylated IgGs. Journal of Allergy and Clinical Immunology, 2012. 129(6): p. 1647–1655.e13.

92. Elsner, R.A., C.J. Hastey, and N. Baumgarth, CD4+ T cells promote antibody production but not sustained affinity maturation during Borrelia burgdorferi infection. Infect Immun, 2015. 83(1): p. 48–56.

93. Chatterjee, S., et al., Serum N-Glycomics Stratifies Bacteremic Patients Infected with Different Pathogens. Journal of Clinical Medicine, 2021. 10(3): p. 516.

94. Ramirez, J., et al., Antibiotics as Major Disruptors of Gut Microbiota. Frontiers in Cellular and Infection Microbiology, 2020. 10.

95. !!! INVALID CITATION !!! [69].

96. Comunale, M.A., et al., Identification and development of fucosylated glycoproteins as biomarkers of primary hepatocellular carcinoma. J Proteome Res, 2009. 8(2): p. 595–602.

97. Comunale, M.A., et al., Linkage Specific Fucosylation of Alpha-1-Antitrypsin in Liver Cirrhosis and Cancer Patients: Implications for a Biomarker of Hepatocellular Carcinoma. PLoS ONE, 2010. 5(8): p. e12419.

98. Guile, G.R., et al., A Rapid High-Resolution High-Performance Liquid Chromatographic Method for Separating Glycan Mixtures and Analyzing Oligosaccharide Profiles. Analytical Biochemistry, 1996. 240(2): p. 210–226.

99. Powers, T.W., et al., MALDI Imaging Mass Spectrometry Profiling of N-Glycans in Formalin-Fixed Paraffin Embedded Clinical Tissue Blocks and Tissue Microarrays. PLoS ONE, 2014. 9(9): p. e106255.

100. Harvey, D.J., et al., Proposal for a standard system for drawing structural diagrams of *N*. PROTEOMICS, 2009. 9(15): p. 3796–3801.

